# Micromanipulation of prophase I chromosomes from mouse spermatocytes reveals high stiffness and gel-like chromatin organization

**DOI:** 10.1101/2020.08.31.276402

**Authors:** Ronald Biggs, Ning Liu, Yiheng Peng, John F. Marko, Huanyu Qiao

## Abstract

Meiosis produces four haploid cells after two successive divisions in sexually reproducing organisms. A critical event during meiosis is construction of the synaptonemal complex (SC), a large, protein-based bridge that physically links homologous chromosomes. The SC facilitates meiotic recombination, chromosome compaction, and the eventual separation of homologous chromosomes at metaphase I. We present experiments directly measuring physical properties of captured mammalian meiotic prophase I chromosomes. Mouse meiotic chromosomes are about ten-fold stiffer than somatic mitotic chromosomes, even for genetic mutants lacking SYCP1, the central element of the SC. Meiotic chromosomes dissolve when treated with nucleases, but only weaken when treated with proteases, suggesting that the SC is not rigidly connected, and that meiotic prophase I chromosomes are a gel meshwork of chromatin, similar to mitotic chromosomes. These results are consistent with a liquid- or liquid-crystal SC, but with SC-chromatin stiff enough to mechanically drive crossover interference.

## Introduction

**M**itosis and meiosis are two forms of cell division that separate chromosomes, but whose purposes and mechanisms are distinct^1^. During mitosis, replicated sister chromatids segregate at anaphase. At the end of mitosis, two daughter cells are formed which are genetically identical to each other and to their mother cell. By contrast, during meiosis, two homologous chromosomes pair up during meiotic prophase I, undergo genetic recombination, and then segregate at anaphase I. The replicated sister chromatids within each of the two homologous chromosomes remain paired until they are separated during meiosis II. The final products of the two rounds of meiotic division are four daughter cells with different genetic makeup. The central events of meiotic prophase I, pairing and synapsis of homologous chromosomes, make meiosis structurally and functionally distinct from mitosis.

The structures of mitotic and meiotic chromosomes are distinct. One of the key differences between mitosis and meiosis is the construction of the synaptonemal complex (SC) during meiotic prophase I ^2, 3, 4^. The SC is a large, tripartite, protein-based lattice, which may act as a mechanically rigid structure that facilitates recombination of homologous replicated chromosomes ^2, 3, 5, 6, 7^. As one goes through the substages of meiosis prophase I, the axial elements of the SC are initially assembled during leptotene, initial bridging (or synapsis) of homologous chromosomes occurs during zygotene, synapsis is completed during pachytene, and finally desynapsis occurs during diplotene. At diplotene, the SC only partially remains, connecting homologous chromosomes at the centromeres and at the chiasmata, the points of genetic crossover^1, 2, 8, 9, 10, 11^. The initial construction of the SC is facilitated by the axial elements SYnaptonemal Complex Protein 2 and 3 (SYCP2 and 3), while the bridging is facilitated by SYCP1 binding to itself and the axial elements^2, 6, 12^. The highly structured SC, its components, and the underlying chromatin have been shown to be highly dynamic^13, 14, 15, 16^, with the SC observed to have liquid-crystalline properties^13, 14, 15, 16^.

The exchange of genetic material during meiosis I is a prerequisite for the proper separation of homologous chromosomes at the end of meiosis I^13, 15^. Meiotic chromosomes appear to limit the number of crossovers, and crossovers localize non-randomly at a set distance if there is more than one along a chromosome^17, 18, 19, 20, 21^. This limitation is known as crossover interference (CI). While there is still much debate concerning the mechanism of CI, one hypothesis suggests that the chromosome contains distinct “beam” and “film” domains, with chromatin dynamics along the beam causing stress fractures in the film allowing the homologous chromosomes to cross over^18, 20^. In this model, the fracture in the film also causes stress reduction in the beam, limiting where another stress fracture/crossover can occur, thus driving CI ^18, 20^.

The protein complexes that act to reorganize chromosomes during mitotic prophase and meiotic prophase I are distinct. The condensin Structural Maintenance of Chromosome (SMC) complex appears to be responsible for much of the morphology and mechanical stiffness of mitotic chromosomes^1, 22, 23^. The cohesin SMC complex is also present on chromosomes during mitosis, generating sister chromatid cohesion, but it is removed from chromosome arms in mitotic prometaphase, remaining predominately at the centromere^24, 25^. In contrast, meiotic prophase I chromosomes contain much less condensin, which does not play a role until later in the meiotic cell division process^5, 10, 26^. Unlike mitosis, meiotic prophase I chromosomes are organized by cohesin, which participates in sister chromatid cohesion, homologous chromosome pairing, construction of the SC, and crossover formation^9, 10, 11, 26, 27^. This basic structure is known as the cohesin core. The lateral elements of the SC are assembled on top of the cohesin core^9, 11, 27^.

Since mitotic prometaphase and meiotic prophase I chromosomes differ structurally, and since the bending stiffness of meiotic chromosomes has been proposed to play a role in CI ^18, 20^, we were motivated to study their structure and mechanics. To do this we developed a technique to physically capture and manipulate meiotic chromosomes to directly study connectivity and mechanical properties relevant to their hypothesized dynamic nature and mechanobiological function (by connectivity we mean the dependence of chromosome structure on disruption of particular molecules, *e.g.*, DNA or specific proteins; by mechanics we primarily mean the elastic response to stretching and bending deformations). This approach has been previously used to investigate newt mitotic chromosomes, human mitotic chromosomes, and female mouse meiotic II chromosomes^22, 28, 29, 30, 31^.

Here we report experiments where we capture and manipulate meiotic chromosomes from mouse meiotic prophase I nuclei, and perform tests on their structure and mechanics, along the lines of prior work on mitotic chromosomes. Due to the fundamental role of SYCP1 in the maturation of the SC and its ability to bridge homologues with synapsis, we also aimed to test if there was a mechanical difference between meiotic chromosomes from WT and null mutant *Sycp1*^−/−^ spermatocytes^32^. Our results show that meiotic prophase chromosomes are much stiffer (ten-fold larger spring constant, normalized to length) than their mitotic counterparts, consistent with the notion that their mechanical stiffness may play a role in CI^19, 20^. As is the case for mitotic chromosomes^30, 31^, meiotic chromosomes behave as “chromatin gels”, in that they can be completely dissolved by nucleases, but only weakened by proteases. This suggests that the SC is not by itself the solid load-bearing structure generating its high stiffness. We discuss how this result is consistent with recent observations of liquid crystal SC behavior^16^.

## Results

### Isolation of meiotic prophase I chromosomes

Previous work on isolated prometaphase mitotic chromosomes revealed novel physical and structural properties, most notably that they are a gel meshwork of chromatin, whose stiffness is dependent on condensin^30, 31, 33^. We utilized a similar approach to isolate meiotic prophase I chromosomes from mouse spermatocytes to study the mechanics of meiotic prophase I chromosomes and identify how SYCP1 contributes to their stiffness. Isolating meiotic prophase I chromosomes was complicated by the presence of the nuclear envelope (Fig. 1a), which is absent during mitotic prometaphase (Fig. 1b). Since meiotic chromosomes’ ends are attached to the nuclear envelope^27, 34^, one end of a chromosome was easily identified and aspirated into a pipette then moved away from the nucleus. The location of the other chromosome end was determined by looking for a change in chromosome morphology (density along the chromosome); that locus was then captured by a second pipette (Fig. 1a). This was followed by removal of the nucleus using a third pipette, which was used to stabilize the nucleus (Fig. 1a).

**Figure 1.**
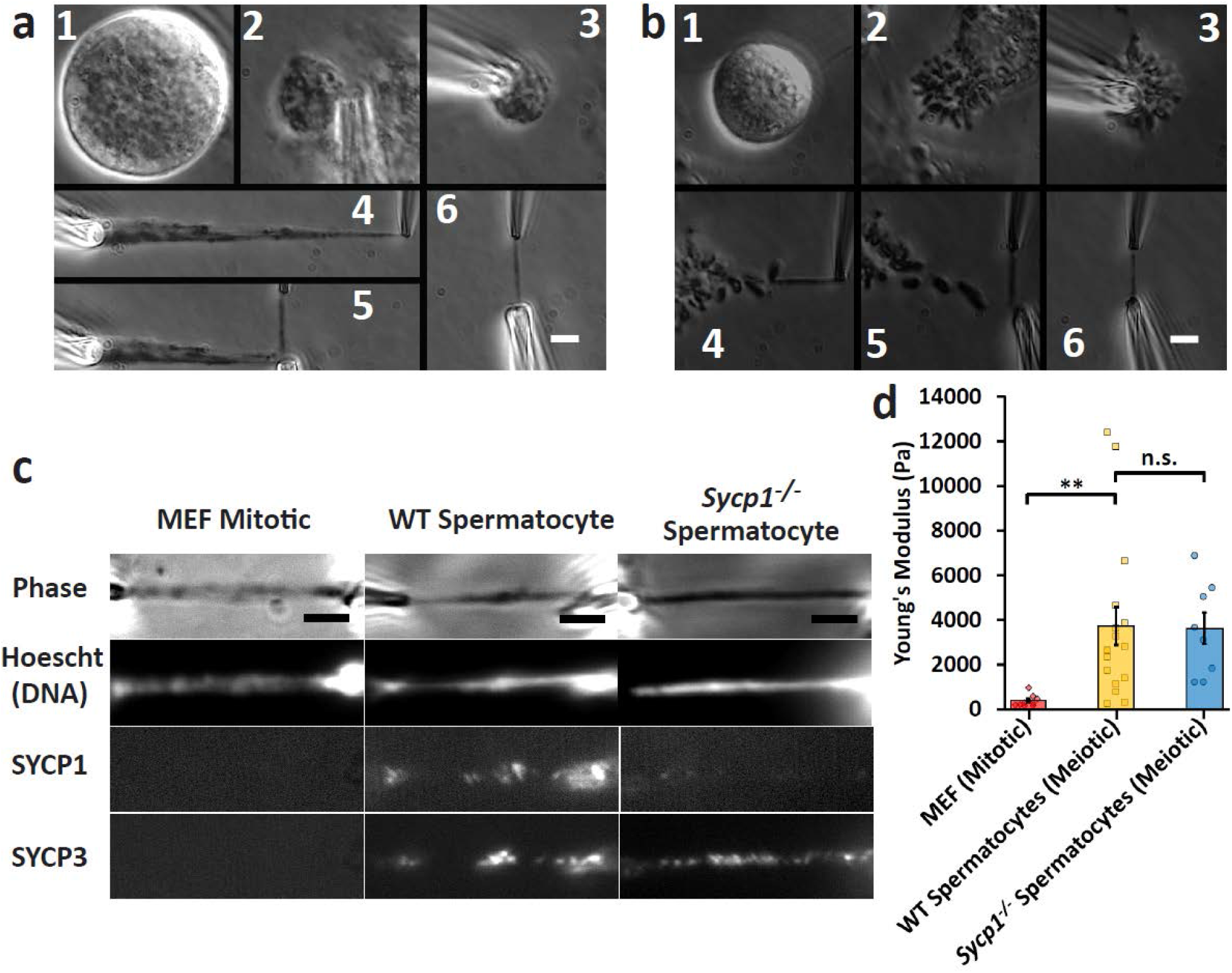
Chromosome isolation and micromanipulation. Images show steps for isolation of meiotic (**a**) and mitotic **(b)** chromosomes. **(a)** A large spermatocyte containing chromosomes (subpanel **1**) was identified, and lysed with Triton-100X in a pipette to release the prophase nucleus **(2)**. Another pipette then was used to grab and hold the nucleus **(3)**. A chromosome end at the edge of the nucleus then was grabbed with a soft force-measuring pipette **(4)**; the other chromosome end was captured with a stiff pulling pipette **(5)**, and the rest of the nucleus was removed, leaving only the prophase chromosomes on the pipettes **(6)**. (**b**) A rounded mitotic cell with visible dark chromosomes in prometaphase was identified **(1)**, lysed with Triton-100X sprayed from a pipette to release the prometaphase mitotic bundle **(2)**, and another pipette then grabbed and held the bundle **(3)**. A chromosome end at the edge of the bundle was grabbed with a thin, force pipette and moved away from the bundle **(4)**, where the other end of the chromosome was grabbed by a stiff, pulling pipette **(5)**, and the bundle was removed, leaving one chromosome isolated between two pipettes **(6)**. Scale bars are 5 μm. (**c**) Distinguishing mitotic MEF from meiotic prophase spermatocyte chromosomes using fluorescence microscopy. MEF chromosomes appear as rod-like structures in the phase-contrast channel that stain for DNA, but do not immunostain for SYCP1 or SYCP3. WT spermatocyte chromosomes appear as rod-like structures that stain for DNA, SYCP1, and SYCP3. *Sycp1*^−/−^ spermatocyte chromosomes stain for DNA and SYCP3, but not SYCP1. Scale bar is 5 μm. (**d**) Stiffness (Young’s Modulus) of mitotic and meiotic chromosomes. MEF chromosomes have a stiffness of 370 ± 70 Pascals (Pa, N=10), ten-fold weaker than WT spermatocyte chromosomes (3700 ± 800 Pa, N=17, *p*=0.006 relative to MEF). WT and *Sycp1*^−/−^ spermatocyte chromosomes (*Sycp1*^−/−^ modulus 3600 ± 700 Pa, N=8, *p*=0.0001 relative to MEF) do not have a statistically significant difference in stiffness (*p*=0.93). All averages are reported as mean value ± SEM. All *p* values calculated via *t* test.

We verified that isolated chromosomes were in meiotic prophase I by immunostaining against SYCP1 and SYCP3 (central and lateral SC components, respectively), which are present only during prophase I of meiosis (Fig. 1c, Supplementary Fig. 1) and which are absent on mitotic chromosomes (Fig. 1c).

### Meiotic prophase I chromosomes are stiffer than mitotic ones

Once isolated, a chromosome could be physically manipulated to measure its Young’s modulus, a measurement of stiffness (see Materials and Methods for more details). Mitotic chromosomes from MEF culture cells have a Young’s modulus of 370 ± 70 Pa, while meiotic chromosomes from wild-type spermatocytes have a modulus of 3670 ± 840 Pa, almost 10-fold stiffer than the mitotic chromosomes (Fig. 1d). In order to determine the effect of the SYCP1 on the stiffness of meiotic chromosomes, we measured the Young’s modulus of prophase I chromosomes from genetic *Sycp1*^−/−^ null mutants^27^. These chromosomes stained positive for DNA and SYCP3, verifying their meiotic prophase I substage, but not SYCP1, showing their lack of a critical SC protein (Fig. 1c, Supplementary Fig. 1). The *Sycp1*^−/−^ chromosomes had a Young’s modulus of 3550 ± 700 Pa, nearly the same as the wild-type spermatocyte chromosomes (Fig. 1d). We note the remarkable difference between the stiffness measurements of Young’s modulus for meiotic and mitotic chromosomes. This suggests that a fundamental substage-independent structure of meiotic chromosomes makes them much stiffer compared to the mitotic chromosomes, despite mitotic chromosomes being more compact. Fig. 2a shows representative images of native (untreated after isolation and capture) chromosome stiffness measurements.

**Figure 2.**
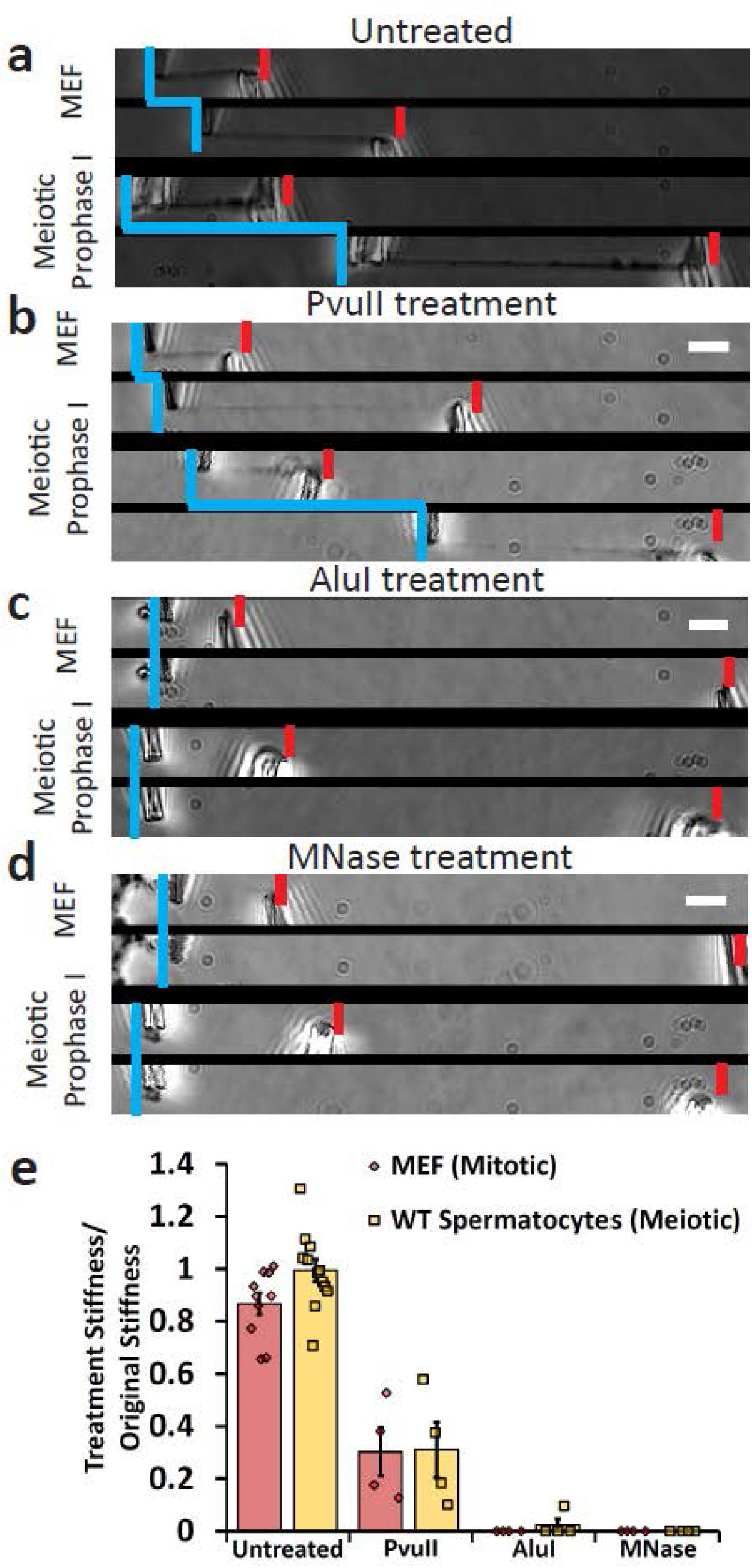
Mitotic and meiotic chromosomes have a contiguous DNA connection, which is dissolved by 4 bp restriction enzymes, but only weakened by 6 bp restriction enzymes. Image pairs show pipette positions untreated (native isolated) chromosomes when relaxed and stretched **(a)**, and chromosomes following enzyme treatments **(b-d)**. Vertical blue lines mark positions of force pipettes. Force pipette deflection by pulling (horizontal blue lines) indicates mechanical connection; no movement (no horizontal blue line) indicates no mechanical connection. Red notches mark positions of stiff pipettes. Bars are 5 μm. (**b**) Both mitotic and meiotic chromosomes were weakened, but not fully digested after treatment with PvuII (cut sequence CAG^˅^CTG). (**c**) Both mitotic and meiotic chromosomes lost connectivity after treatment with AluI (cut sequence AG^˅^CT; for 1 of 4 trials meiotic chromosomes were not fully digested by AluI). (**d**) Both mitotic and meiotic chromosomes lost connectivity when treated with MNase (cleaves all DNA sequences). (**e**) Quantification of chromosome stretching elasticity after no treatment or after being treated with PvuII, AluI, and MNase. No treatment caused a 13 ± 4% weakening of mitotic chromosomes (N=10) and a 1 ± 4% weakening of meiotic chromosomes (N=10). PvuII treatment caused a 70 ± 8% reduction in stiffness for MEF chromosomes (N=4) and 70 ± 9% reduction in stiffness for meiotic chromosomes (N=4). One of four AluI treatments of meiotic chromosomes caused a 90% reduction in stiffness (rather than fully digesting), while AluI treatment digested 4 of 4 mitotic chromosomes. All MNase treatments caused full digestion of mitotic and meiotic chromosomes (N=4 in both cases). All averages are reported as mean value ± SEM. Bars are 5 μm.

### Nucleases dissolve meiotic prophase I chromosomes

In order to probe the underlying structure of the isolated meiotic chromosomes, we treated them with proteases and nucleases^30, 31^. Mitotic chromosomes were merely weakened when treated with a 6-bp-sequence-specific restriction enzyme (PvuII, Fig. 2b), but completely dissolved (disconnected) when treated with 4-base pair (bp) sequence-specific (AluI, Fig. 2c) and non-sequence-specific (MNase, Fig. 2d) nucleases ^30, 31^. Meiotic prophase I spermatocyte chromosomes behave nearly identically to mitotic chromosomes when treated with nucleases (Fig. 2b-d). MNase treatment also removed all connectivity between mitotic bundles and meiotic prophase nuclei (Supplementary Fig. 2).

### Proteases do not dissolve meiotic prophase I chromosomes

Treatment with the proteases trypsin and proteinase K only caused mitotic MEF chromosomes to weaken, not disintegrate (Fig. 3a-c). Meiotic prophase I chromosomes were also not fully disintegrated when treated with both proteases and were less affected by protease treatment compared to mitotic chromosomes (Fig. 3a-c). This we found remarkable considering the amount of additional protein structures present on meiotic chromosomes, emphasizing the importance of chromatin (*i.e.*, DNA) connectivity to their organization and mechanics.

**Figure 3.**
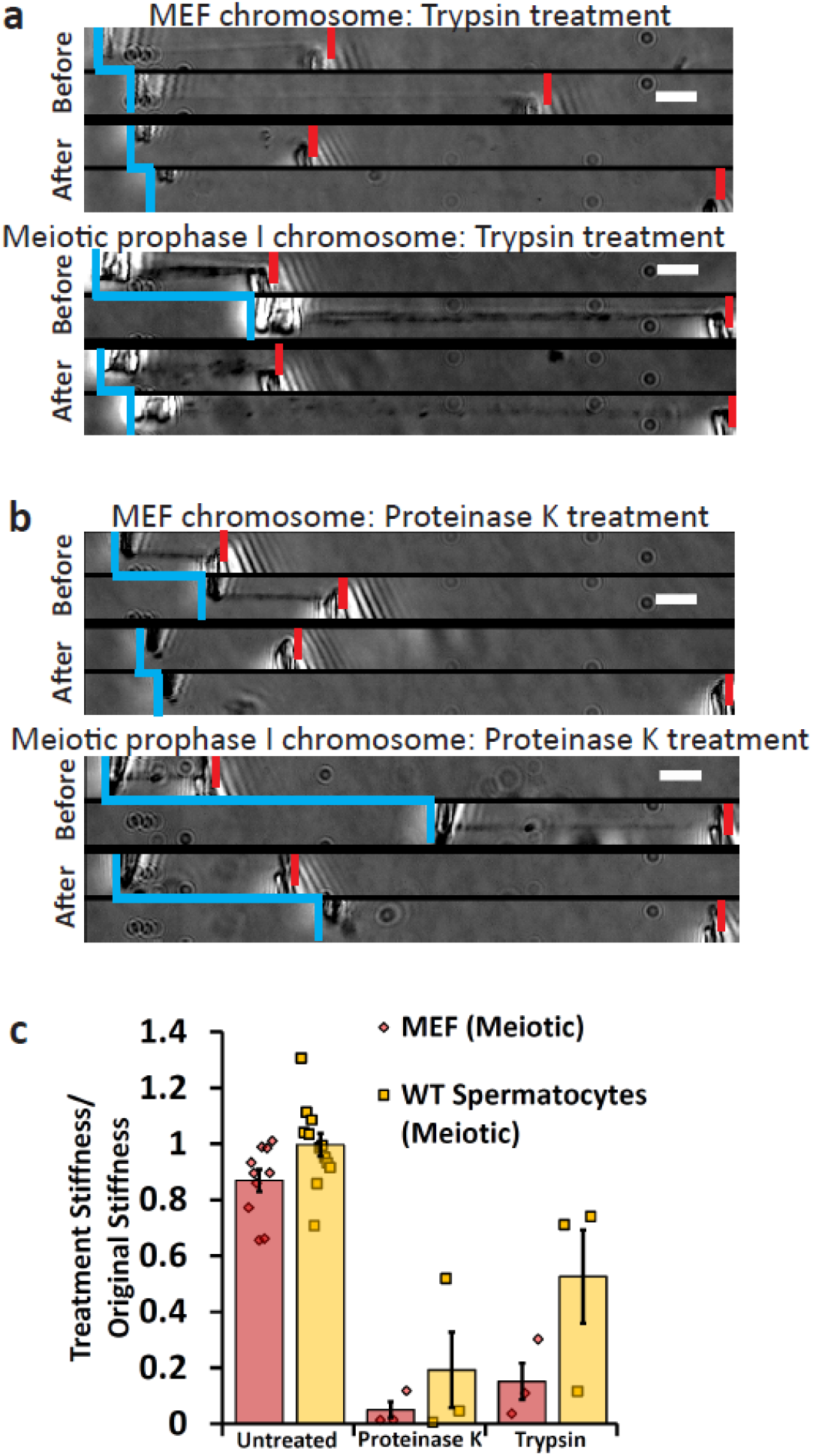
Both meiotic and mitotic chromosomes are weakened, but do not dissolve when treated with Trypsin and Proteinase K. **(a, b)** Vertical blue lines indicate force-measuring pipette positions. Force pipette movement means connection (indicated by horizontal blue line connecting relaxed and stretched force pipette positions); no movement means no connection. Red notches mark positions of stiff pipettes. Image pairs show pipettes for relaxed and stretched chromosomes. Before enzyme treatments, the left pipette is deflected indicating force transmitted through the chromosome. Bars are 5 μm. **(a)** MEF chromosomes lose all phase contrast after treatment with trypsin; but still are well connected enough to move the force pipette, and are weaker than before treatment (smaller deflection during pulling, After panel). Similarly, meiotic chromosomes lose most definition in the phase-contrast channel, but can move force pipette after digestion when treated with Trypsin, and are weaker than before treatment. **(b)** MEF chromosomes lose all definition in the phase-contrast channel, but can move force pipette after digestion when treated with Proteinase K and are weaker than before treatment. Similarly, meiotic chromosomes lose most definition in the phase-contrast channel, but can move force pipette after digestion by Proteinase K and are weaker than before treatment. **(c)** Quantification of untreated and protease-treatment weakening of mitotic and meiotic chromosomes. No treatment caused a 13 ± 4% weakening of mitotic chromosomes (N=10) and a 1 ± 4% weakening of meiotic chromosomes (N=10). Trypsin treatment caused an 85 ± 6% weakening in mitotic chromosomes (N=3) and a 48 ± 17% weakening in meiotic chromosomes (N=3). Proteinase K treatment caused a 95 ± 3% weakening in meiotic chromosomes (N=3) and an 81 ± 13% weakening in meiotic chromosomes (N=3). All averages are reported as mean value ± SEM.

## Discussion

By capturing and manipulating meiotic prophase I chromosomes and comparing them to mitotic chromosomes, several interesting similarities and differences in their structure are revealed. First, both meiotic prophase I and mitotic chromosomes show the same underlying structure, *i.e.* a chromatin gel meshwork crosslinked by protein complexes (possibly predominantly SMC complexes^22, 26, 30^). This general model is supported by experiments where both types of chromosome are fully disintegrated when treated with nucleases but maintain their connectivity (remain mechanically connected between pipettes) when treated with proteases, albeit while losing most of their stiffness. Meiotic cohesin is present between the arms of sister chromatids through meiotic prophase I^9^. Likewise, condensin can be found compacting mitotic chromosomes along their arms^22, 23^. If these complexes formed a contiguous structure through self-attachment or other forms of protein bridging, then the contiguous protein structure would remain between the pipettes after nuclease treatment^30^. This digestion occurs in the wild-type spermatocytes, indicating that the presence of SYCP1 is not sufficient to prevent this digestion.

We note that although we suspect most of the meiotic prophase chromosomes that we have studied are in pachytene (synapsed), based on observation of SYCP1 staining along their lengths and a single-filament morphology, we have not yet established this definitively. We do not yet have protocols for meiotic substage identification that are compatible with our capture and micromanipulation experiments. It is possible that the chromosomes we have studied are not fully synapsed, and that the fully synapsed SC forms a contiguous structure that is resistant to nuclease digestion^6^. Further experiments could be done to test this hypothesis: a result where meiotic chromosomes with intact SCs dissolved during nuclease treatment in pachytene (where the SC is fully synapsed) would support a highly dynamic (*e.g.*, liquid crystal^16^) SC organization.

It is also possible that isolation of the chromosomes to outside of the cell combined with the dynamic nature of the SC leads to the SC developing a non-contiguous structure when isolated, for example as a result of a liquid-like SC^16^. Gaps in the protein network, connected by chromatin, would therefore dissolve with nuclease treatment. These gaps would leave behind fragments of the SC, carried away by the flow of the spray liquid. This would eventually lead to the entire dissolution of the meiotic chromosome. These different scenarios are subjects for further study, but we emphasize that we have found that straightforward isolation of meiotic chromosomes led reproducibly and systematically to objects that were dissolved by nucleases, indicating a strong dependence on chromatin itself for their structural integrity.

By measuring their Young’s moduli, derived from stretching experiments, we found meiotic prophase I chromosomes are 10-fold stiffer than mitotic prometaphase chromosomes, demonstrating their differences. The origin of this difference could be due to the SC, both in terms of its components and construction during meiotic prophase I^7, 18^. Although meiotic prophase I chromosomes’ stiffness is similar to the stiffness of meiotic II arrested chromosomes^28^. The underlying chromatin structure could also impact meiotic chromosome stiffness, as could meiotic bivalents, the resolution of chromatin entanglements, crossovers, and other such differences^7, 15, 18, 27, 35, 36^. Another factor that may underlie the high stiffness is the difference between the resolution and compaction of chromosomes over time, leading to a strong difference between mechanics of prophase and prometaphase chromosomes. Comparison of mitotic prophase and meiotic prophase chromosome mechanics would be of interest in this regard but as yet we have been unable to isolate mitotic prophase chromosomes for mechanical study, due to the difficulty of extracting them from somatic cell nuclei.

The high Young’s modulus of meiotic prophase I chromosomes has a potentially important implication for CI. The bending of a beam can be thought of as differential stretching across its cross-section, indicating that the difference in stretching elasticity of meiotic prophase and mitotic chromosomes that we have observed predicts a similar difference in bending stiffnesses^37^. Therefore, the function of a large stiffness of meiotic prophase chromosomes is qualitatively consistent with the stress model of CI, which requires a large elastic stiffness to allow a single crossover event to affect a meiotic chromosome along its entire length^18, 20^. This higher stiffness, not required or observed for mitotic chromosomes, could originate in the underlying chromatin being more tightly packed and/or crosslinked in meiotic prophase I (*e.g.,* by meiotic cohesin and SC components)^6, 9, 16, 26^. Determining the quantitative dependence of degree of CI and meiotic chromosome stiffness, for example using knockdowns or knockouts of specific proteins involved in CI, could provide tests for the stress-release model. While there are no theoretical studies predicting the quantitative stiffness needed to drive CI in the mechanical stress model, our measurements provide a starting point for quantitative experimental tests of it.

While the meiotic chromosome is stiff, its stiffness did not depend strongly on the presence of SYCP1, the central element of the SC. This was quite surprising, since the presence of a central element might be expected to act as a crosslinker of material across homologous chromosomes^6, 7, 14, 27^. This crosslinking would bridge homologue pairs together and presumably stiffen the whole structure. However, it is possible that this finding is again consistent with the idea that the SC is very dynamic *in vivo*^15, 16^. This dynamic *in vivo* nature could lead to the SC only weakly contributing to the overall mechanical stiffness of the meiotic chromosome, although this may be due to its isolation from the intracellular environment. Future investigations into the SC include the role of synapsis of SYCP1 across meiotic substages and the roles of SYCP2 and SYCP3^2, 11, 12^. The underlying cohesin core of meiotic chromosomes promises to be an exciting topic for further investigation^6, 15, 26, 27, 34^, since when mitotic chromosomes lack condensin, the main mitotic SMC complex, they completely lose their morphology and stiffness^23^. Disruption of the cohesin SMC complex might be expected to drive similar strong effects during meiotic prophase. We also note that direct measurement of bending stiffness of meiotic chromosomes would be desirable to verify the expected relation between their stretching and bending elasticities^38, 39^.

The complex nature and dynamic structure of meiotic chromosomes are important to understand because of their essential role in sexual reproduction. Being physical objects, studying their mechanical properties furthers our understanding of how they function in the cell (*e.g.*, in determining the mechanism underlying CI)^18^. Further experiments of the general type introduced here, comparing stiffness and nuclease sensitivity across the substages of meiotic prophase may reveal the underlying mechanical function of the SC and its role in the connectivity of meiotic prophase I chromosomes. Mechanical experiments targeting the lateral elements of the SC could determine the difference in mechanical makeup of the SC by focusing on the elements loaded first onto the SC^9, 10, 26^. Isolation and manipulation of cohesin deficient mutants could facilitate discovery of the underlying crosslinkers of meiotic chromosomes, the impact of meiotic SMC complexes, and the contribution of sister chromatid cohesion in meiosis to the chromosomal structure^27^. Other functions such as the sensitivity of mechanics to alteration of topoisomerase presence or activity are also likely to uncover interesting relationships between chromatin entanglements and stiffness^18, 21, 31^.

## Methods

### Animals

Male wildtype and *Sycp1*^−/−^ C57BL/6 mice were used in this study^32, 40, 41^. All mice were housed in the Animal Care Facility at the University of Illinois Urbana-Champaign (UIUC) under 12 h dark/12 h light cycles at 22 ± 1°C. Animal handling and procedures were approved by the UIUC Institutional Animal Care and Use Committee. Mice used in the study were adults from 2 to 6 months in age.

### Cell culture and spermatocyte samples

Mouse testes were extracted from adult mice with or without the *Sycp1* null mutation and sent on ice to be dissected for extraction of spermatocytes containing meiotic prophase I cells for chromosome isolation. The testes were kept in PBS or DMEM at 4 °C for up to 3 days, after which the cells would appear damaged and the chromosomes not suitable for isolation. Spermatocyte samples were extracted from the testes by cutting at the surface of the testes (to maximize the number of meiosis I spermatocytes) in ~150 μL PBS and put into a well containing ~1.5 mL PBS for single chromosome isolation.

Viability tests were performed using Trypan Blue on cells kept for 0-4 days. While there is a marked decrease in cells that prohibit the entrance of Trypan Blue over time, indicating viable cells, there are still many cells that exhibit normal Trypan Blue exclusion (Supplementary Fig. 3). In our experiments, we verified that the cell had a normal and intact cell membrane phenotype, a rounded cell that dissolved when sprayed with 0.05% Triton X-100-PBS solution. All the cells were imaged for documentation for this visualization, examples shown in Supplementary Fig. 4.

Experiments on mitotic cell used mouse embryonic fibroblast (MEF) cells, which were maintained in DMEM (Corning) with 10% fetal bovine serum (FBS) (HyClone) and 1% 100x penicillin streptomycin (Corning). The cells were incubated at 37 °C and 5% CO_2_ for no more than 30 generations, passaged every 2-4 days. The cells were plated and allowed to recover 1-3 days before chromosome isolation (Fig. 1b). All mitotic cells performed in free-cycling cells and mitotic cells were identified by eye without drugs.

### Isolation and microscopy of meiotic chromosomes

Single chromosome isolation experiments used an inverted microscope (IX-70; Olympus) with a 60x 1.42 NA oil immersion objective with a 1.5x magnification pullout at room temperature and atmospheric CO_2_ levels. Experiments were performed in less than 3 hours after removal from the 4 °C refrigerator (meiotic spermatocytes) or 37 °C incubator (MEF cells) to ensure minimum damage to the cells and chromosomes being analyzed.

Meiotic prophase I (spermatocytes) and prometaphase (MEF) cells were identified by eye and lysed with 0.05% Triton X-100 in PBS (Fig. 1a,b). All other pipettes were filled with PBS. After lysis, the meiotic nucleus or mitotic bundle was held with a pipette. One end of a loose chromosome was grabbed by the force pipette, moved from the nucleus or bundle, and grabbed with the pulling pipette on the other end. The nucleus or bundle was then removed to isolate the chromosome (Fig. 1 a,b).

### Single chromosome force measurements

An easily bent force pipette (WPI TW100F-6), and stiff pulling pipette (WPI TW100-6) were used for stretching chromosomes, forged using a micropipette puller (Sutter P-97) and a custom pipette cutting setup. Once isolated, the pipettes were moved perpendicular to the chromosome, stretching the chromosome to roughly its native length (Fig. 2a). The stiff pipette was then moved 6.0 μm and returned to the starting position at a constant rate of 0.20 μm/sec in 0.04 μm steps in a LabVIEW protocol, while it tracked the stiff and force pipette. Refer to Supplementary Fig. 5 for an example of force-extension trace. The deflection of the force pipette multiplied by its calibrated spring constant then divided by the distance between the pipettes (the stretch) to obtain the chromosome spring constant. The chromosome spring constant multiplied by its initial length gave the doubling force. Cross-sectional area was estimated as πr^2^/2. The diameter was calculated as the full width at half maximum of an ImageJ scan. All other physical measurements taken (chromosomal doubling force, spring constant, initial length, and cross-sectional area) reported and shown in Supplementary Fig. 6.

Young’s (elastic) modulus expresses the stress/strain ratio and is calculated with the formula E=(F/A)/(ΔL/L_0_), where E is the Young’s modulus, F is the force the object is under, A is its cross-sectional area, ΔL is the change in length from L_0_, which is the initial, unstretched length of the object. Rearranged, the Young’s modulus can also be written as E=K*L_0_/A, where K is the spring constant of the object (F/ΔL). Our calculations in bulk are thus: the linear regression of the deflection-stretch line, multiplied by the force pipette spring constant, multiplied by the initial length, and divided by the cross-sectional area of the chromosome. See Supplementary Fig. 5 for representative images for the linear regression line of the force-extension graph.

### Single chromosome immunofluorescence

In immunofluorescence experiments the chromosome was lifted above the glass surface after mechanical measurements, then sprayed with a primary, secondary, and tertiary solution from a wide-bore pipette, moving the chromosome between sprays. The solutions used 50 μL PBS, 36-38 μL H_2_O (Corning), 10 μL 5% casein (Sigma), and 2 μL each antibody. Experiments used a rabbit anti-SYCP1 (GeneTex) with a mouse anti-SYCP3 (Santa Cruz Biotechnology) primary and a 488-nm anti-mouse IgG (Invitrogen) with a 594-nm anti-rabbit IgG (Invitrogen). The tertiary spray used Hoechst instead of an antibody.

### Single chromosome enzyme spray experiments

In enzymatic digestion experiments on chromosomes, the chromosome would be sprayed after force measurements. For experiments on nuclei or chromosome bundles, the nuclei or chromosome bundle would be sprayed directly after isolation. All solutions were diluted to 100 μL with water except proteinase K (which did not need specialized buffer) after adding buffer and enzyme. 10 μL of this solution was then aspirated into a wide-bore pipette and sprayed onto the chromosome, nuclei, or chromosome bundle for around 5-10 min. The number in parentheses refers to the final concentration of enzyme. MNase (non-specific DNA digestion) used MNase buffer and 2 μL of enzyme (20 units/μL) PvuII (NEB), AluI (NEB) (cut sequence AG^˅^CT) used NEBuffer 2.1 and 10 μL of enzyme (1 unit/μL), PvuII (cut sequence CAG^˅^CTG) used NEBuffer 3.1 and 10 μL of enzyme (1 unit/μL), Trypsin (NEB) (cuts after Lys/Arg) used 2x Trypsin buffer and 50 μL of Trypsin diluted in 200 μL water (50 ng/μL), Proteinase K (NEB) (digests most peptide bonds) used PBS as the buffer and used 2 μL (200 ng/μL).

### Statistical Methods and Reproducibility

Means, standard deviations, and standard errors were computed for sets of data using standard office software. Experiments were carried out approximately 10 times and *p* values were computed using the t-test. Experimental measurements were carried out using different cells (replicates), with the entire data sets derived from at least three separate animals’ tissue.

## Acknowledgements

We thank Supipi Mirihagalle, Lois Suh, and Tianming You for help with mouse husbandry and data collection. Work at NU was supported by NIH grants R01-GM105847, U54-CA193419 (CR-PS-OC) and a subcontract to grant U54-DK107980 (4D Nucleome). Work at University of Illinois, Urbana-Champaign was supported by NIH grants R00-HD082375 and R01-GM135549.

## Author contributions

R.B., J.F.M. and H.Q. conceived and designed the research. R.B., N.L., Y.P. and H.Q. performed the experiments. R.B., N.L., Y.P. and H.Q. analyzed the results. R.B., N.L., Y.P., J.F.M. and H.Q. wrote the manuscript.

## Data availability

All measurements are included in the accompanying Supplementary Materials. Additional data (e.g., raw, uncalibrated instrument data files) are available from the corresponding author upon request.

