## Supplementary Figures for "Micromanipulation of prophase I chromosomes from mouse spermatocytes reveals high stiffness and gel-like chromatin organization"

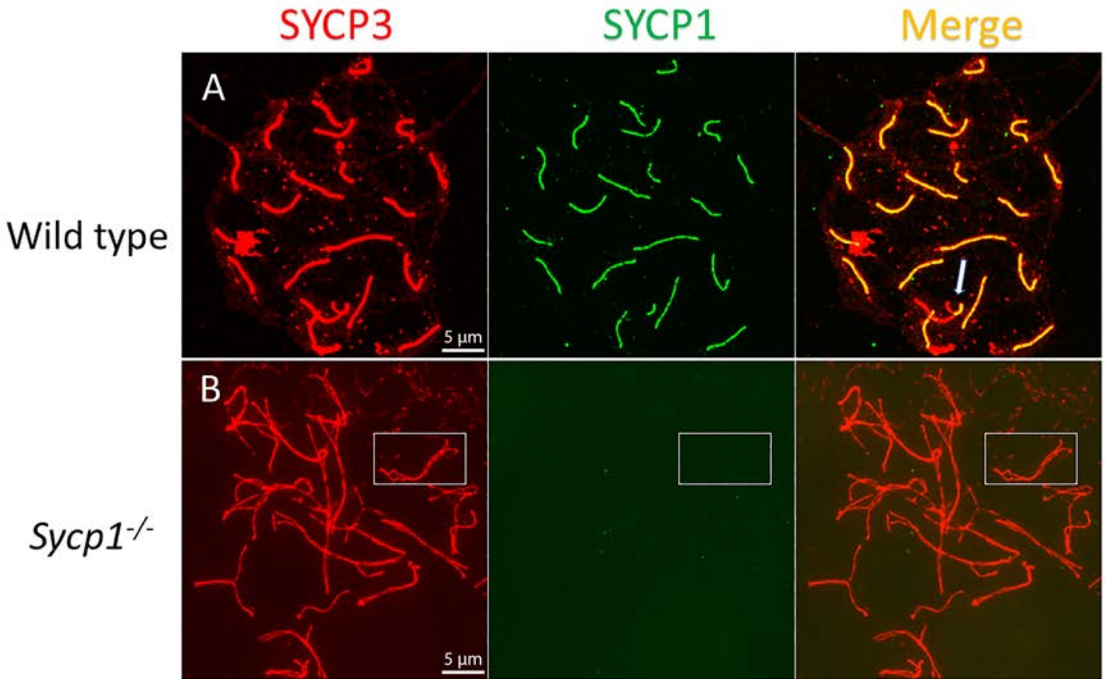


**Supplementary Figure 1. SYCP1 is absent in *Sycp1* knockout mutant spermatocytes.** Immunostaining for SYCP3 (red) and SYCP1 (green) in wild-type (A) and *Sycp1*$^{-/-}$ (B) spermatocytes. (A) In wild-type pachytene spermatocytes, all homologous chromosome pairs synapsed from one end to the other except sex chromosomes. Synapsis only occurs in the Pseudo Autosomal Regions (PAR) of the sex chromosome pairs (arrow). (B) Synaptonemal complexes do not form between the paired homologous chromosomes in *Sycp1*$^{-/-}$. One homolog pair is highlighted by a white frame. Note that there are no transverse filaments, SYCP1, between the two homologous chromosomes.

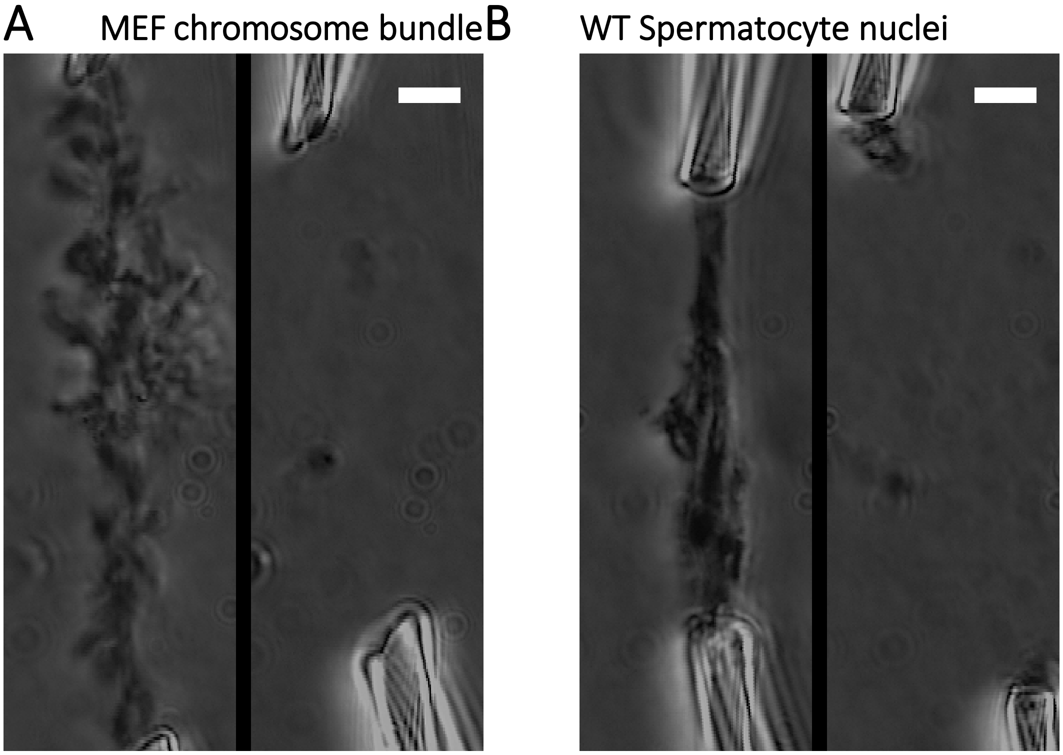


**Supplementary Figure 2. Example MNase treatment of captured mitotic bundles and meiotic nuclei, both stretched between two pipettes.** (A) MNase treatment of mitotic bundles causes loss of any chromosomes as seen in phase-contrast imaging. (B) MNase treatment of meiotic nuclei causes loss of any chromosomes as seen in phase-contrast imaging.

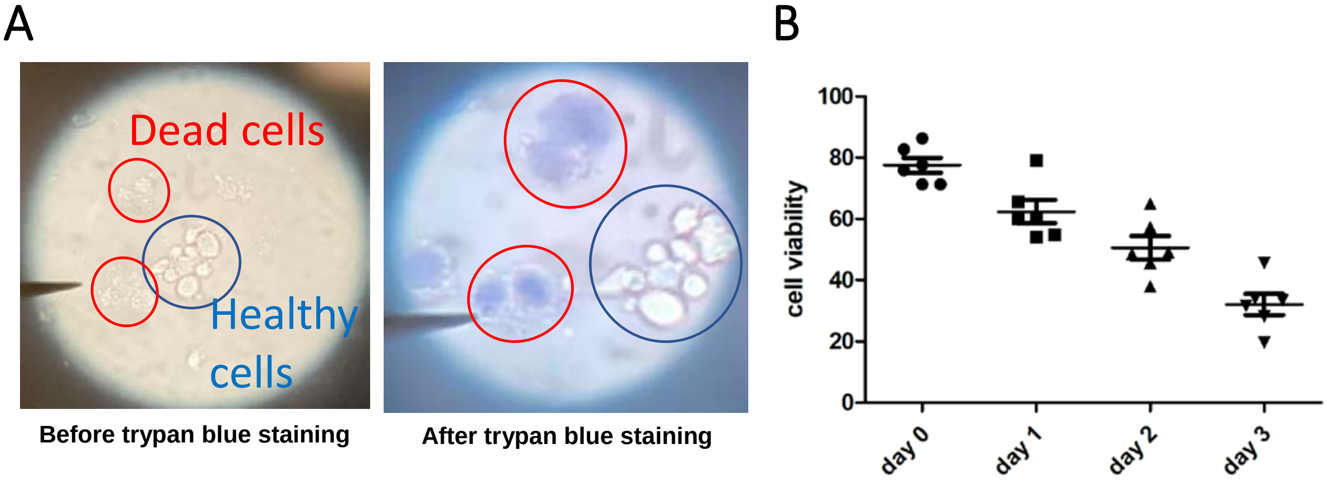


**Supplementary Figure 3. Trypan blue cell viability assay.** (A) Trypan blue stain invades dead cells without an intact cell membrane but is excluded from living cells. Brightfield image of the field of view is corresponding with the trypan blue imaging to demonstrate difference of living and dead cells. (B) Quantification of spermatocyte cell viability over time, start point of day testes extracted.

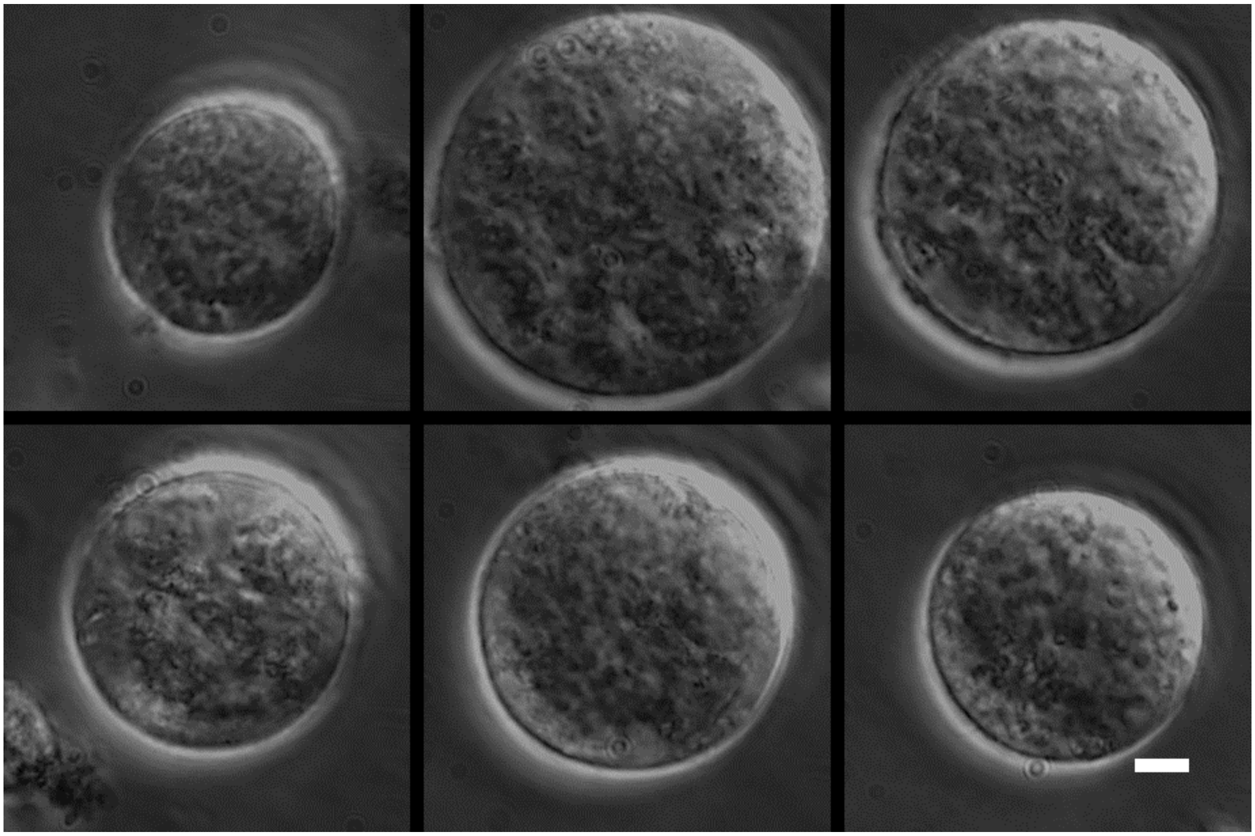

**Supplementary Figure 4. Example images of wild-type (*Sycp1*$^{+/+}$) meiotic cells before nuclei extraction.** Example of the *Sycp1*$^{+/+}$ meiotic cells with intact cell membrane before isolation where the chromosomes were extracted (Scale bar = 5μm)

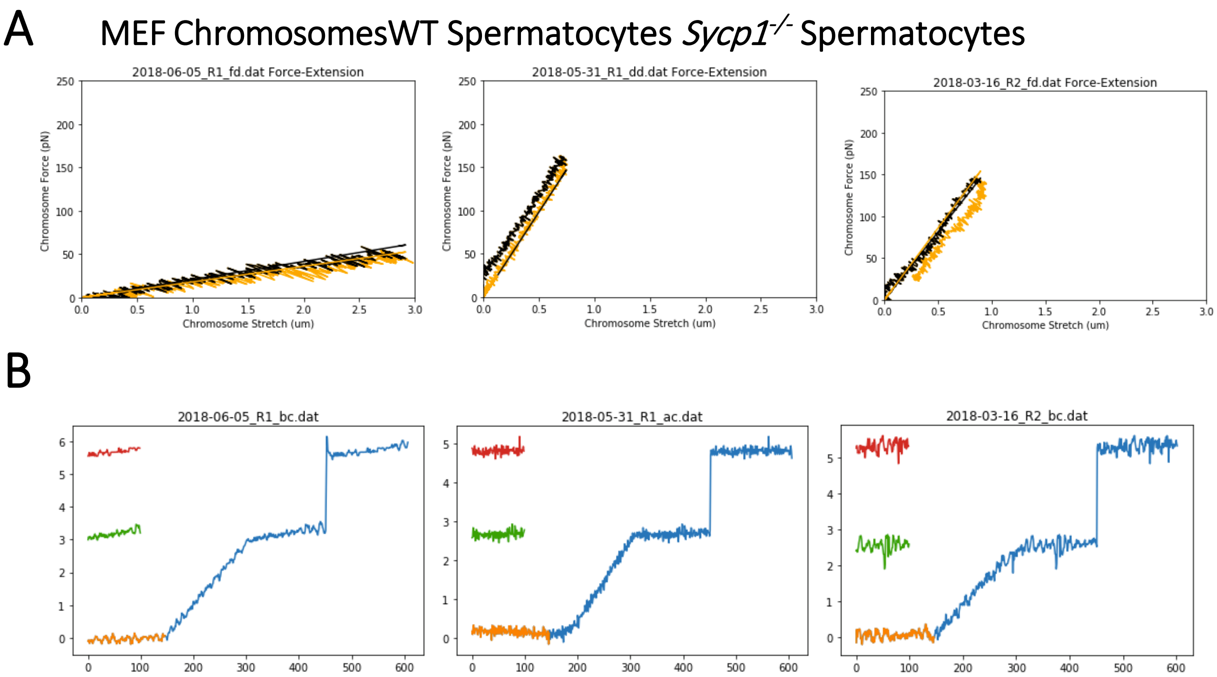


**Supplementary Figure 5. Example Force-Extension plots and corresponding calibration plots.** (A) Representative Force-Extension plot for mitotic MEF, meiotic wild-type, and *Sycp1*$^{-/-}$ mutant chromosomes, respectively (left to right). The black lines trace the outgoing trace and linear regression slope, while the orange lines track the return trace and return linear regression slope. (B) Example trace of a force pipette calibration against the same calibration standard pipette. The orange trace shows the original position of the pipette. The green trace shows the position when the calibration pipette was moved but held back by the unknown force pipette stiffness. The red trace shows the full deflection of the calibration pipette. The blue trace shows the position of the calibration pipette over the entire run.

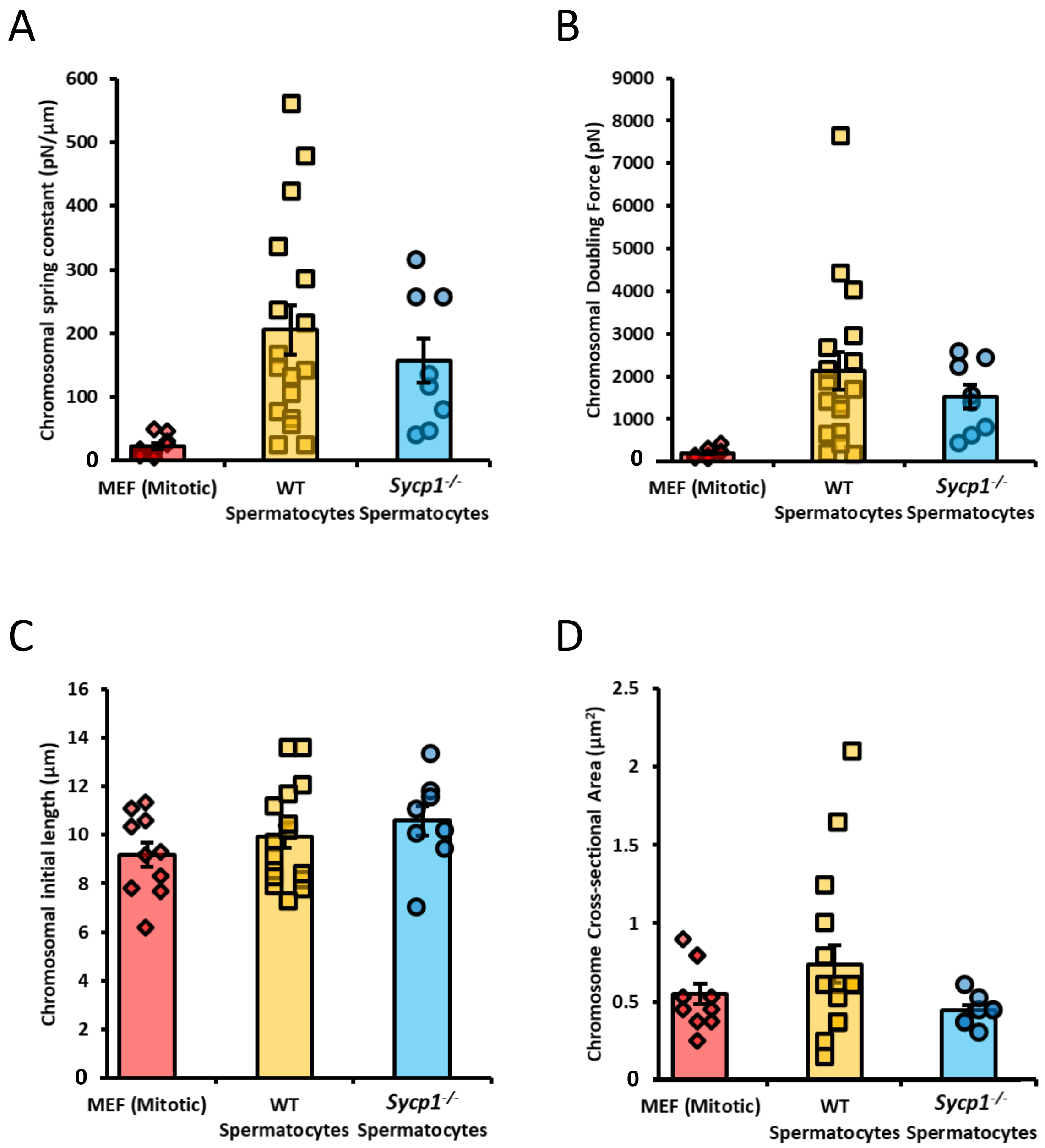


**Supplementary Figure 6. Additional physical measurements of mitotic and meiotic chromosomes.** (A) Spring constant distribution of chromosomes (the force it takes to stretch a chromosome a micron). Chromosomes from MEF culture cells have a spring constant of 22 ± 5 pN/µm, WT Spermatocytes of 206 ± 38 pN/µm, *Sycp1*$^{-/-}$ spermatocytes 157 ± 35 pN/µm. MEF chromosomes were statistically significantly different from the spermatocytes ($p$=0.004 for MEF *vs.* WT; $p$=0.0009 for MEF *vs Sycp1*$^{-/-}$), while the spermatocytes were not significantly different from each other ($p$=0.56 for WT *vs* *Sycp1*$^{-/-}$). (B) Doubling force distribution of chromosomes (the force required to stretch the chromosome twice its initial length). Chromosomes from MEF culture cells have a doubling force of 190 ± 40 pN, WT Spermatocytes of 2130 ± 440 pN, *Sycp1*$^{-/-}$ spermatocytes 1520 ± 280 pN. MEF chromosomes were statistically significantly different from the spermatocytes ($p$=0.008 for MEF *vs.* WT; $p$=0.001 for MEF *vs Sycp1*$^{-/-}$), while the results for spermatocytes were not significantly different from each other ($p$=0.51 for WT *vs* *Sycp1*$^{-/-}$). (C). Initial length distribution of chromosomes (the length of the chromosome unstretched/at rest/isolated from the cell). Chromosomes from MEF culture cells have an initial length of 9.2 ± 0.5 µm, WT Spermatocytes of 9.9 ± 0.5 µm, *Sycp1*$^{-/-}$ spermatocytes 10.6 ± 0.6 µm. All groups were not significantly different from each other ($p$= 0.43 for MEF *vs* WT spermatocytes, $p$= 0.11 for MEF *vs* *Sycp1*$^{-/-}$ spermatocytes, and $p$=0.37 for WT *vs* *Sycp1*$^{-/-}$ spermatocytes). (D) Cross sectional area distribution of chromosomes (the area of the chromosome thickness, estimated as the area of a cylinder using the full width at half maximum derived from a line scan in ImageJ). Chromosomes from MEF culture cells have an area of 0.55 ± 0.06 µm$^2$, WT Spermatocytes of 0.74 ± 0.12 µm$^2$, *Sycp1*$^{-/-}$ spermatocytes 0.44 ± 0.03 µm$^2$. All groups were not significantly different from each other ($p$=0.17 for MEF vs WT spermatocytes, $p$=0.18 for MEF *vs* *Sycp1*$^{-/-}$ spermatocytes, and $p$=0.057 for WT spermatocytes *vs* *Sycp1*$^{-/-}$ spermatocytes). All averages reported as mean value ± SEM. All $p$ values calculated via $t$ test.
